# Simultaneous global labeling (SiGL) of 5-methylcytosine and 5-hydroxymethylcytosine by DNA alkylation with a synthetic cofactor and engineered methyltransferase

**DOI:** 10.1101/2022.10.31.513813

**Authors:** Sigal Avraham, Leonie Schütz, Larissa Käver, Andreas Dankers, Sapir Margalit, Yael Michaeli, Shahar Zirkin, Dmitry Torchinsky, Noa Gilat, Omer Bahr, Gil Nifker, Maya Koren-Michowitz, Elmar Weinhold, Yuval Ebenstein

**Affiliations:** Department of Chemistry, Raymond and Beverly SacklerFaculty of Exact Sciences, Department of Biomedical Engineering, Tel Aviv University, 6997801Tel Aviv, Israel; School of Chemistry, Tel Aviv University, Ramat Aviv, Tel Aviv 6997801 (Israel); Institute of Organic Chemistry, RWTH Aachen University, D-52056 Aachen Germany 3; Hematology department, Shamir medical center, Be’er Ya’akov Israel

**Keywords:** 5-hydroxymethylcytosine, DNA methylation, DNA methyltransferase, epigenetic biomarker, fluorescence

## Abstract

5-methylcytosine and 5-hydroxymethylcytosine are epigenetic modifications involved in gene regulation and cancer. Here, we describe a new, simple, and high-throughput platform for multi-colour epigenetic analysis. The novelty of our approach is the ability to multiplex methylation and de-methylation signals in the same assay. We utilize an engineered methyltransferase enzyme that recognizes and labels all unmodified CpG sites with a fluorescent cofactor. In combination with the already established labelling of the de-methylation mark 5-hydroxymethylcytosine via enzymatic glycosylation, we obtained a robust platform for simultaneous epigenetic analysis of these marks. We assessed the global epigenetic levels in multiple samples of colorectal cancer and observed a reduction in 5-hydroxymethylcytosine levels, but no change in DNA methylation levels between sick and healthy individuals. We also measured epigenetic modifications in chronic lymphocytic leukaemia and observed a decrease in both modification levels. Our results indicate that this assay may be used for the epigenetic characterization of clinical samples for research and patient management.

## Introduction

5-methylcytosine (5mC) is created by the addition of a methyl group to carbon at the 5 position of cytosine by methyltransferase enzymes (DNMTs or MTases)^1^. These enzymes are responsible for the generation and maintenance of genomic methylation patterns^2^, which play a crucial role in various cellular processes^3,4^. In mammals, DNA methylation occurs mostly on cytosine of CG sequence motifs, commonly referred to as CpGs^3^. The methyl group is donated from *S*-adenosyl-L-methionine (SAM or AdoMet), a universal cofactor involved in methyl group transfer^5,6^. DNA methylation is a stable and heritable DNA modification, but it can be reversed in an active DNA demethylation process, where in the first step, 5mC is oxidized to 5-hydroxymethylcytosine (5hmC) by the ten-eleven translocation (TET) family of dioxygenases^7^. Since its rediscovery in 2009, 5hmC has become the focus of many studies, and it has been found that 5hmC is involved in physiological processes and cancer^8,9^. While global 5hmC levels decrease in most cancers ^9–13^, 5mC has been shown to increase or decrease depending on the cancer type. Importantly, the commonly used bisulfite treatment for analysis of DNA methylation does not distinguish between 5mC and 5hmC and thus reports a convolution of both levels. Global epigenetic analysis of both 5mC and 5hmC can serve in several key areas, such as early cancer detection, monitoring of disease progression, and response to treatment^9^.

Various approaches have been developed and are widely used for global detection and quantitation of 5mC and 5hmC. The most sensitive method for profiling cytosine DNA methylation is LC-MS/MS. Although this method is accurate and requires relatively small amounts (50-100 ng) of DNA to analyze, it requires expertise, expensive equipment, and demanding assay optimization^14,15^. Simultaneous monitoring of global 5mC and 5hmC is possible with liquid chromatography-electrospray ionization tandem mass spectrometry with multiple reaction monitoring (LC–ESI–MS/MS–MRM). This method is fast, robust, and accurate; however, it requires expertise and cannot monitor multiple samples at once^16^. Other available methods are DNA dot-blot^8,17–19^, immunohistochemically (IHC) staining^20–24^, and enzyme-linked immunosorbent assays (ELISA)^25–28^. Although these assays are relatively fast and straightforward, they tend to produce high error rates^29^ and lack the sensitivity required to track minute but significant epigenetic content changes crucial for identifying related medical conditions^11^. Furthermore, these assays are not compatible with simultaneous detection of both 5mC and 5hmC due to the bulkiness of the used antibodies and contrast agents. To overcome these limitations, we developed a new technique that utilizes custom-made multi-sample array slides that enable sensitive quantification of a large number of samples quickly and accurately, opening an avenue for large-scale DNA epigenetic modification monitoring for research and clinical use^9^.

Unlike 5hmC which is amenable to direct optical detection in genomic DNA samples^9,11,30–32^, direct labelling of cytosine methylation is extremely challenging due to the inert nature of the methyl group. An alternative is to label unmodified cytosines by enzymatic alkylation. The quantity of unmodified CpGs (um-CpGs) is inversely correlated to genomic methylation levels. DNA MTase enzymes may utilize synthetic cofactor analogs as alkylation agents in order to transfer functional groups to DNA^33–36^. CpG methyltransferases are blocked when the cytosine is methylated, and thus will label only um-CpGs (Figure 2a). We and others have shown that a double mutant of the CpG-specific DNA MTase M.Sssl can label unmodified cytosines with azide groups *via* the AdoYnAzide cofactor^37,38^. However, M.Sssl is difficult to express at high concentrations and it tends to aggregate upon expression. Here, we introduce a corresponding double mutant of the homologous CpG-specific DNA MTase M.MpeI from the bacterium *Mycoplasma penetrans* for rapid and efficient labelling of um-CpGs.

The double mutant (dm) M.MpeI is easy to produce at high concentrations and may serve as a robust alkylation agent for methylation detection with AdoMet analogues. The enzyme transfers azide groups to unmodified cytosines in CpG sites, and then a fluorophore is clicked on by strain-promoted azide-alkyne cycloaddition (SPAAC) for optical detection (Figure 1a). DNA samples are deposited on multi-sample array slides^9,39^ and fluorescence intensity is read-out as an inverse measure of DNA methylation (Figure 1b). Here, we develop and validate the M.MpeI (dm) assay for 5mC analysis and then combine it with direct 5hmC labelling using two different colours for simultaneous detection of both epigenetic marks. We also successfully apply the combined assay to colon cancer and chronic lymphocytic leukaemia (CLL) samples.

**Figure 1.**
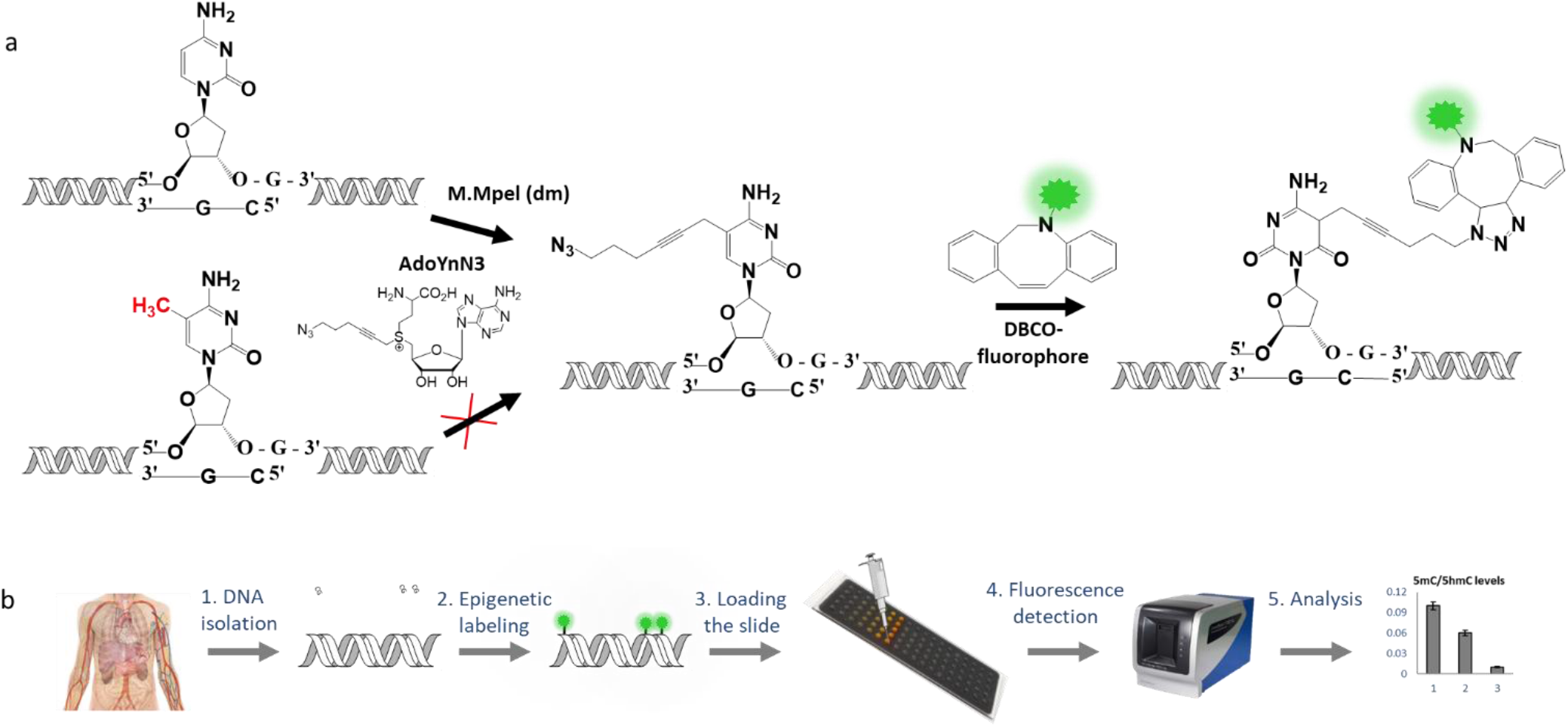
**a)** Schematic illustration of the two-step chemoenzymatic fluorescent labelling of um-CpG sites with the Mmpel (dm) enzyme. **b)** Schematic illustration of the slide assay’s workflow. 1) DNA extraction from the tissue of interest. 2) Two-step fluorescent labelling reactions for 5hmC residues/ um-CpG sites and sample purification from excess fluorophores. 3) Deposition of labelled DNA samples on the activated multi-sample array slides. 4) Fluorescence imaging in a commercial slide scanner. 5) Data analysis and 5hmC/um-CpG quantification.

## Results and Discussion

M.MpeI (dm) is CpG specific and efficiently transfers azides to DNA. The wild type M.MpeI (Figures 2a and 2b) has a sterically obstructed active site and is only capable of using AdoMet as a cofactor (Figure S1). Therefore, the active site was enlarged by site-directed mutagenesis of amino acids Q136 and N374 to A (Figure 2c), leading to an M.MpeI double mutant (dm). To assess the activity of the mutant it was challenged with several AdoMet analogues with extended methyl group replacements (Table S1) and screened using a modification/restriction assay. Figure 3 shows that M.MpeI WT is only active with the natural cofactor AdoMet while M.MpeI Q136A/N374A (M.MpeI dm) shows also activity with AdoYnYn (lower, lane 7) and AdoYnAzide (lower, lane 9).

**Figure 2.**
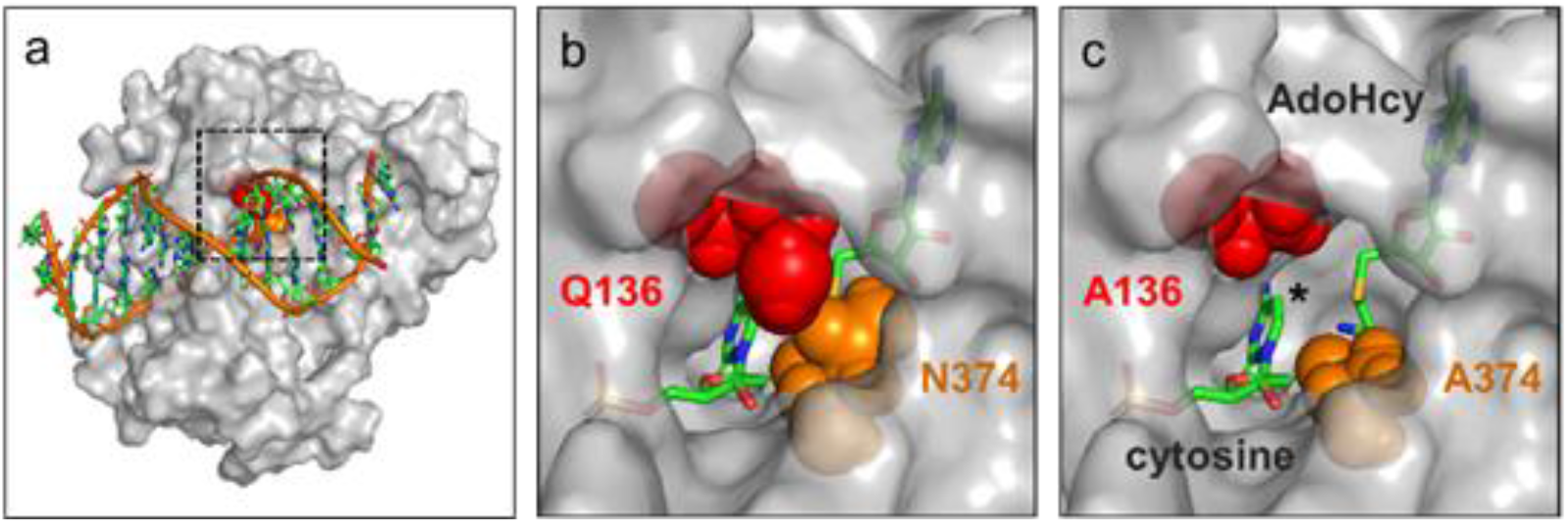
Structure of the DNA cytosine-C5 MTase M.MpeI (grey surface representations) in complex with DNA (ribbon) and the cofactor product *S*-adenosyl-L-homocysteine (AdoHcy) (PDB ID: 4dkj). **a)** Overall structure of the complex with the boxed active site. **b)** The active site of the wild-type enzyme is obstructed by amino acids Q136 and N374. **c)** Mutations to A136 and A374 are expected to enlarge the active site to accommodate cofactor analogs with extended methyl group replacements for direct transfer of lager alkyl groups from the sulfur atom (yellow) of the cofactor to C5 of the target cytosine (indicated by a star).

**Figure 3.**
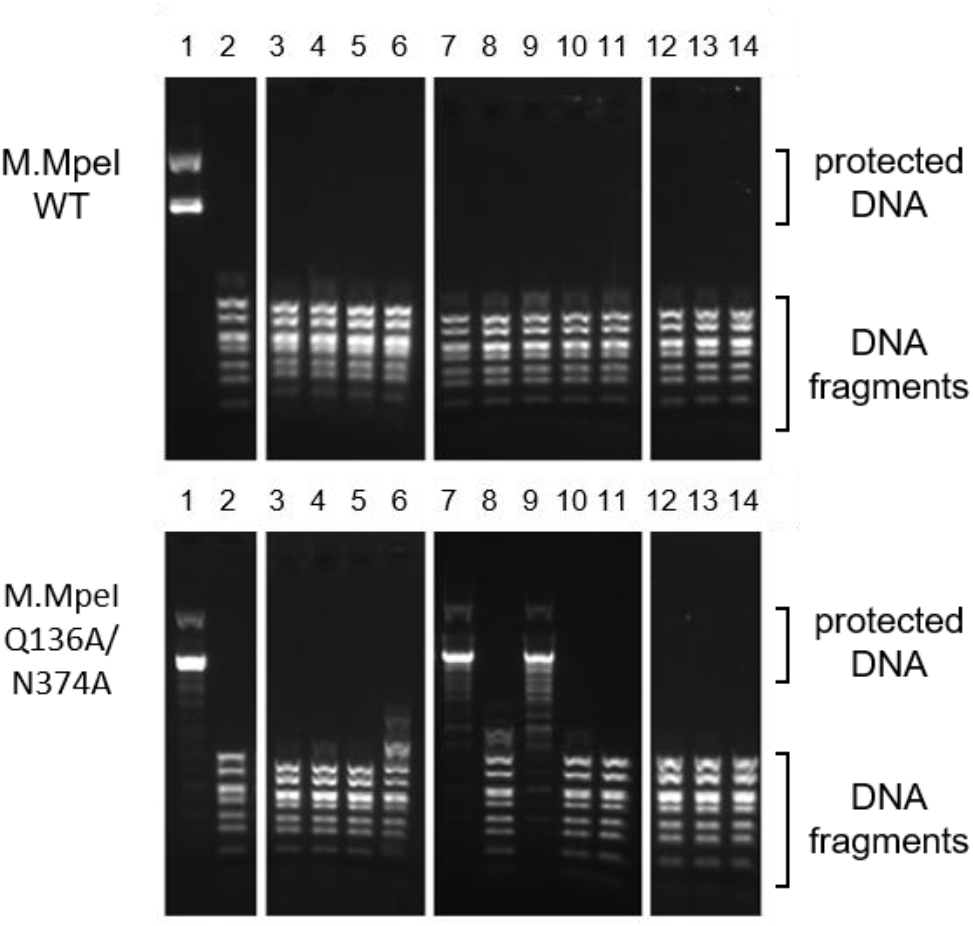
Cofactor screening by modification/restriction assay. Plasmid DNA is incubated with M.MpeI wild type (WT) (upper) or with M.MpeI Q136A/N374A variant (lower) and AdoMet (lane 1), no cofactor (lane 2) or AdoMet analogues with extended methyl group replacements (lanes 3–14). Afterward, the DNA is challenged with the modification-sensitive restriction endonuclease R.BstUI (5’-CGCG-3’). Active MTase/cofactor combinations result in DNA protection against cleavage by R.BstUI while inactive combinations lead to DNA fragmentation.

In order to verify that M.MpeI (dm) labels CpG dinucleotides specifically, we performed reverse-phase HPLC (RP-HPLC) of nucleosides obtained after enzymatic fragmentation of DNA treated with M.MpeI (dm) and AdoYnAzide. As seen in figure 4 and the rustling calculation, M.MpeI (dm) specifically labels CpG dinucleotides and no other cytosine residue.

**Figure 4.**
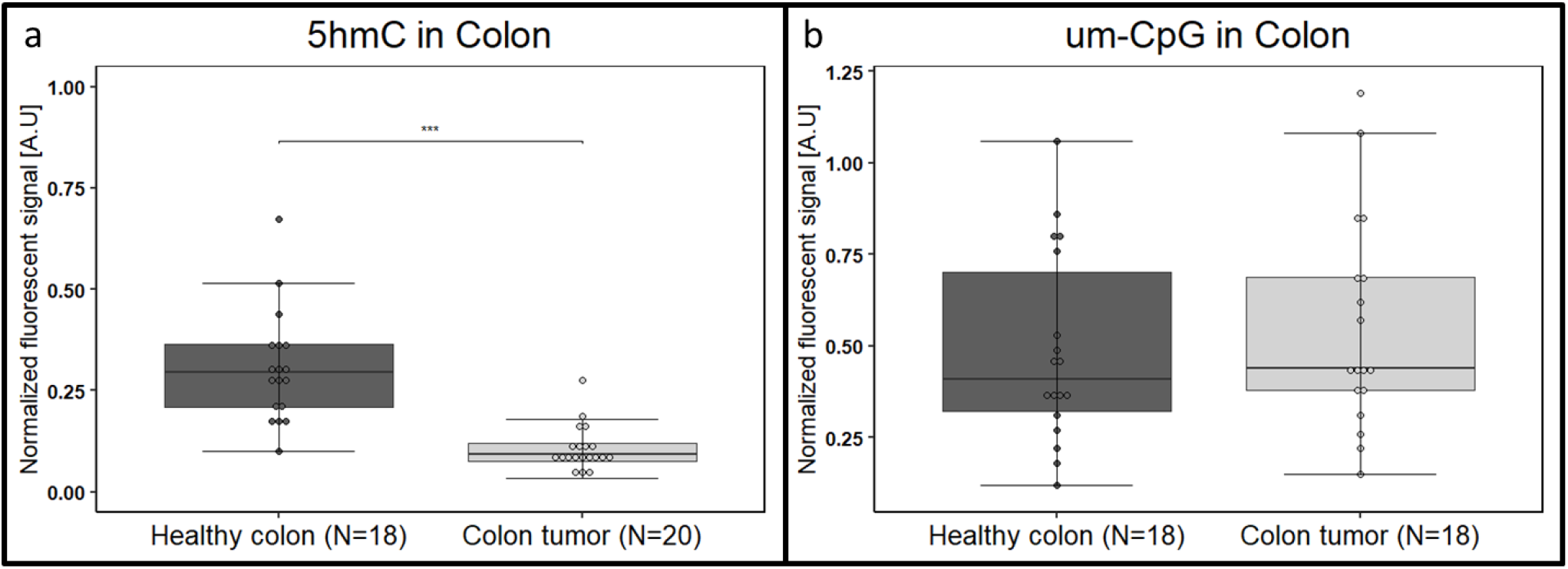
Boxplot representation of colon samples. The boxes correspond to the 25–75 percentiles of the data. The line inside the boxes stands for the median. The whiskers include values within 1.5 times the interquartile range **a)** Relative fluorescent intensity of 5hmC in colon samples. Boxplot representation of 5hmC level in the healthy colon (N=18) and colon tumour samples (N = 20). p < 0.001. **b)** Relative fluorescent intensity of um-CpG labelling, which represents the methylation level in colon samples. Boxplot represents 5mC level in healthy colon (N = 18) and colon tumour samples (N = 18). p >0.1.

**Table 1.**
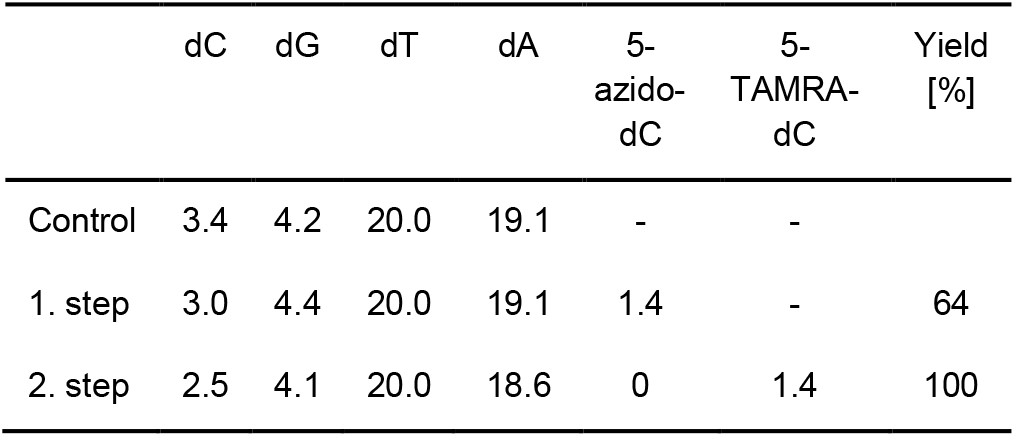
Quantification of CpG labelling with M.MpeI Q136A/N374A by RP-HPLC analysis (see Figure 4). Amounts of nucleosides obtained after duplex modification with M.MpeI Q136A/N374A and AdoYnAzide (1. step) followed by click labelling with DBCO-PEG4-TAMRA (2. step).

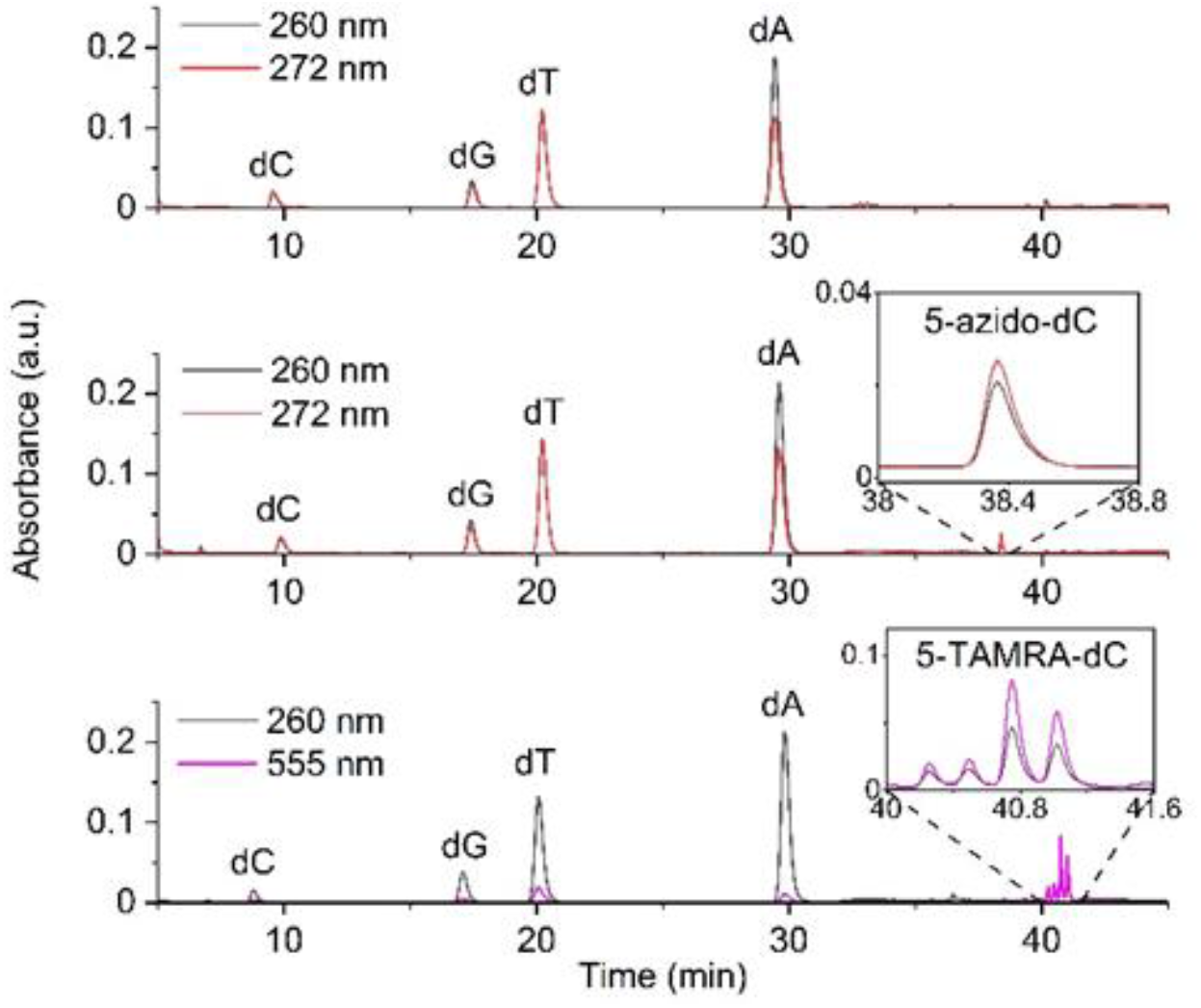

### Assay validation

Several validation measures were carried out in order to design a multi-sample array platform for rapid quantification of 5mC and 5hmC (see supporting information figures S3-S6). In short, we first verified that the assay fluorescence readout is linear with respect to both DNA fragment length and the number of available CpGs. The signal remained linear for both green (TAMRA) and red (Cy5) fluorophores. Next, we characterized the limit of detection (LOD) in order to determine the minimal level of methylation detectable in a given sample.

We calculated a LOD of 0.0024% modification to total nucleotides in a typical sample of 25ng genomic DNA (figure S4). This LOD is lower than the 5hmC content in blood, the tissue with the lowest levels of 5hmC in the human body. Moreover, the LOD value is even lower than that measured for the blood cancer patients, highlighting the assay’s sensitivity and ability to reliably measure and distinguish between healthy and sick individuals even at these low 5hmC levels. Nevertheless, LOD may be increased by the deposition of larger amounts of DNA per sample.

In terms of labelling efficiency, the slide assay displayed an efficiency of 15% and 12% for the TAMRA and Cy5 fluorophores, respectively (figure S6). Although this labelling efficiency is lower than that observed for ODN by RP-HPLC, it is sufficient for reliable results using the slide assay for global detection and quantification of um-CpG.

### Colon cancer

The following results for 5hmC labelling in colon samples were previously published by our group and these measurements are now extended to um-CpG as well^9^.

To assess the performance of the assay for analysis of colorectal samples, we measured 5hmC levels in samples from healthy human colons compared to CRC tumours. We observed a significant reduction in 5hmC for CRC tissue relative to healthy colon (0.108±0.055, N = 20 vs. 0.306±0.139, N = 18, p < 0.001; Figure 4.a).

Then we labelled the same colon samples for um-CpGs using the M.MpeI (dm) labelling reaction. The results show a large variance between patients in each group. In contrast to 5hmC results, the average between cancer patients and healthy ones stays approximately the same (2.82±1.85, N = 18 vs. 2.40±1.46, N = 18, p >0.01; Figure. 4.b), implying there was no detectible change in global methylation between sick and healthy individuals. The reason may be that in the case of methylation, both decrease and increase are observed in cancer, depending on the type and stage of the disease. These results emphasize the importance of investigating the two types of modifications in parallel for cancer research.

### Simultaneous global labelling: blood cancer – CLL

After establishing the two labelling reactions for 5hmC and um-CpGs separately, we performed them on the same DNA sample to get more layers of information in one experiment and demonstrate the clinical relevance of global quantification of um-CpGs and 5hmC using this assay. DNA from whole blood and PBMCs of CLL patients and healthy individuals was fluorescently labelled for 5hmC residues using DBCO-Cy5 and after cleaning from excess fluorophores, labelled for um-CpGs using DBCO-TAMRA (figure 5.a). The labelled DNA was imaged on a single-molecule fluorescence microscope (TILL Photonics) to verify the co-labelling of both epigenetic modifications on the same DNA molecule. The DNA’s backbone was stained with the intercalating dye YOYO-1 while the um-CpGs and 5hmC labels were detected along the DNA molecules as fluorescent spots in different colours. As shown in the representative images in figure 5.b the DNA molecules exhibit 5hmC and um-CpG labelling.

**Figure 5.**
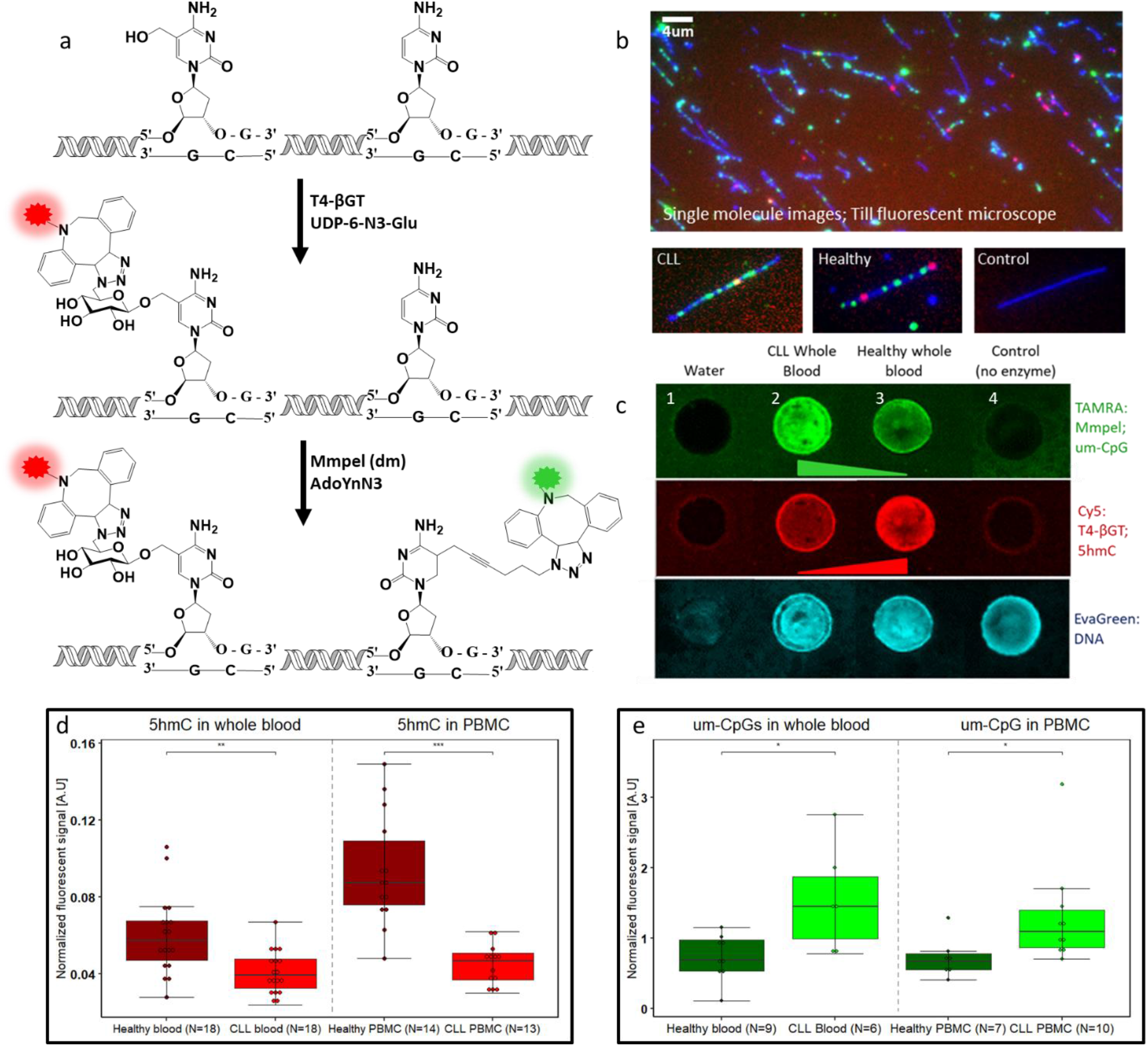
Simultaneous global labelling in hematological samples - SiGL. **a)** Schematic illustration of 5hmC and um-CpG labelling. First, 5hmC residuals were labelled using DBCO-CY5 fluorophore, and then the um-CpG sites were labelled with the new Mmpel (dm) enzyme using DBCO-TAMRA. **b)** Full field of view (left) of human DNA stained with YOYO-1 intercalator dye (blue). 5hmC residues are labelled with DBCO-Cy5 (red), and um-CpG sites are labelled with DBCO-TAMRA (green). Enlarged DNA molecules cropped from various fields of view (bottom); CLL whole blood (left), healthy whole blood (middle), and a control sample (right). **c)** Representative slide image of um-CpG labelling (green) and 5hmC labelling (red) in the blood of a CLL patient (2) and healthy whole blood (3). Column 1 contains double distilled water (DDW) and serves as a control for the DNA channel (cyan). Column 4 is a control sample, performed without the addition of the respective enzyme. **d)** Boxplot representation of 5hmC labelling in whole blood and PBMC samples from CLL patients compares to healthy individuals (whole blood 0.041±0.011, N = 18 vs. 0.060±0.020, N = 18, p < 0.01; PBMCs 0.045±0.010, N = 13 vs. 0.093±0.029, N = 14, p < 0.001). **e)** Boxplot representation of um-CpG labelling in whole blood and PBMC samples from CLL patients compares to healthy individuals (whole blood 1.545±0.742, N = 6 vs. 0.725±0.318, N = 9, p < 0.05; PBMCs 1.304±0.726, N = 10 vs. 0.719±0.284, N = 7, p < 0.05). All boxes correspond to the 25–75 percentiles of the data. The line inside the boxes stands for the median. The whiskers include values within 1.5 times the interquartile range.

Next, the labelled samples were deposited on a mult-sample arrrray slide and fluorescently imaged on the slide scanner in three colours (figure 5.c). Using these measurements, we quantified the intensities of 5hmC and um-CpG for each tissue. We observed a significant reduction in 5hmC levels in both whole blood and PBMCs from CLL patients when compared to the same tissue type from healthy individuals (whole blood 0.041±0.011, N = 18 vs. 0.060±0.020, N = 18, p < 0.01; PBMCs 0.045±0.010, N = 13 vs. 0.093±0.029, N = 14, p < 0.001; Figure 5.d).

These results suggest that the method is sensitive enough to detect the reduction in 5hmC levels in samples with physiological low 5hmC content. When comparing um-CpG levels between sick and healthy individuals, methylation levels also decreased in the cancerous tissues compared to the healthy ones of the same tissue type (whole blood 1.545±0.742, N = 6 vs. 0.725±0.318, N = 9, p < 0.05; PBMCs 1.304±0.726, N = 10 vs. 0.719±0.284, N = 7, p < 0.05; Figure 5.e).

Furthermore, the similarity between the two different hematological samples for both 5hmC and um-CpG levels proposes that molecular quantification of these epigenetic modifications can be performed from blood, which is the most accessible tissue for diagnostic applications.

## Conclusion

5-methylcytosine (5mC) and 5-hydroxymethylcytosine (5hmC) are epigenetic modifications involved in gene regulation, and their global alteration has been reported in different types of cancer. Global analysis of 5mC and 5hmC can potentially serve as biomarkers for early detection, monitoring disease progression, and responding to treatment. Most current techniques for global quantitation of 5mC and 5hmC lack the sensitivity required for clinical applications while being applicable to one modification type at a time.

This work presents the development of a method for direct global analysis of epigenetic modifications. The novelty of this approach is the utility of a new methyltransferase enzyme, M.MpeI (dm), that recognizes unmodified CpG sites, and in combination with a synthetic cofactor, can couple a fluorophore to unmethylated cytosine. Mmpel improvespreviously reported enzymatic labelling by MTaql, covering only about 5.5%^40^ of CpG sites across the human genome (hg38).

We verified the specificity and linear labelling dependency on methylation content of the M.MpeI (dm) reaction, and compared the labelling to a commercial Methyltransferase enzyme (Figure S4). After establishing the labelling reaction, we combined the M.MpeI (dm) labelling with the glucosylation-based 5hmC labelling and achieved simultaneous labelling of um-CpGs and 5hmC modifications on the same DNA sample, enabling more layers of information in one experiment.

Our assay allows concurrent, sensitive, and high-throughput detection, which can potentially be utilized for clinical cancer diagnostics. This is made possible by the low LOD (0.0024%), which is lower than the 5hmC levels found in blood cancer, and allowed us to distinguish between healthy and sick individuals even at these low 5hmC levels. In combination with custom multi-sample array slides and fluorescence imaging on a commercial slide scanner, we obtained a robust platform for the simultaneous analysis of um-CpG and 5hmC global levels in two important types of cancer. We established the global levels of um-CpGs and 5hmC in over 18 samples of colorectal cancer and saw a 3.5-folds reduction in 5hmC levels, but no change in 5mC between sick and healthy individuals. We also measured epigenetic modifications in blood cancer (CLL) and observed a decrease in both modification levels (5hmC: whole blood 30%; PBMCs 40%. 5mC: whole blood 53%; PBMCs 48%.). Moreover, when comparing the different types of samples (whole blood and PBMC) within the cancer patients, um-CpG and 5hmC levels stay approximately the same, suggesting that a simple blood test may be sufficient to perform the analysis, which greatly eases samplehandling for diagnostic applications.

### Experimental Section

The detailed conditions and protocol for the 5hmC labeling, slide handling, and data analysis appear in the supporting information and were adopted from procedures recently published by our lab^9,39^.

### Um-CpG labelling with M.MpeI (dm)

Um-CpGs were fluorescently labelled via a two-step chemoenzymatic reaction. In each reaction tube, 500 ng of DNA were mixed with 2 μL of 1 mg/mL (X10) BSA (New England Biolabs), 2 μL of 10x Mmpel buffer (50 mM Tris-HCl, 100 mM NaCl, 5% glycerine, pH 7.5) that was autoclaved after preparation, 1.33 μL of 75% glycerol, 2 μL of 100 mM 2-mercaptoethanol to a final concentration of 10 mM, AdoYnN3 to a final concentration of 80 μM, Mmpel (dm) to a final concentration of 5.33 μM and ultrapure water to a final volume of 20 μL. The reaction mixture was incubated for 2 hours at 37°C. Then, 20 μg of Proteinase K (PK) (Sigma) was added and incubated for 1 hour at 55°C. Next, the samples were incubated at 80°C for 20 minutes for heat inactivation. Following inactivation, Dibenzocyclooctyl (DBCO)-PEG4-5/6-TAMRA or Dibenzocyclooctyl (DBCO)-Sulfo-Cy5 (Jena Bioscience, Jena, Germany) was added to a final concentration of 250 μM and then incubated overnight at 37°C. The labelled DNA samples were purified from excess fluorophores using Oligo Clean & Concentrator columns (Zymo Research), according to the manufacturer’s recommendations, with three washing steps and two elutions for optimal results.

### Simultaneous global labelling (SiGL)

The dual-colour labelling reactions were performed sequentially. First, the 5hmC labelling (see SI) was performed, and after cleaning from excess fluorophores (Oligo Clean & Concentrator, Zymo research), the samples were labelled with a different fluorophore for um-CpG and purified (Oligo Clean & Concentrator, Zymo Research) before imaging.

### Multi-sample array slides preparation

Multi-well, epoxy -coated microscope slides covered with a Teflon multi-well mask(Tekdon, customized well formation, 2 mm diameter wells, 90 wells per slide) were immersed in 0.005% poly-L-lysine solution in water (Sigma) in order to positively charge the surface. The immersed slides were incubated for one hour at 37°C with mild shaking (25 rpm) and then incubated overnight at 4°C. The following day a blocking step was performed; the 20 slides were washed twice with PBST (0.05% Tween 20, Sigma) solution and twice with PBS (Sigma) and immersed in a 5% w/v bovine serum albumin (Sigma) solution in PBS. The immersed slides were incubated for one hour at 37°C with mild shaking (25 rpm) and then incubated overnight at 4°C. In the final step, slides were washed with water and dried under a flow of nitrogen gas. The slides were used immediately upon drying.

### DNA attachment to the activated slides

1 μL of labelled-DNA samples was placed in each well. The optimal DNA concentration for attachment is 10-30 ng per well. From each sample, 3-5 replicates were placed on the slide. Slides were incubated for 14 minutes at 42°C and then for 24 minutes at 30°C, in humid conditions to avoid rapid drying of the wells (Thermoshaker, Eppendorf). The slides were then washed with water and dried under a flow of nitrogen gas. To avoid light exposure, slides were kept in the dark.

### DNA staining

Total DNA was stained with EvaGreen DNA binding dye (Biotium). 1 μl of 1.25 μM dye (90% water, 10% DMSO) was added to the wells containing the bound DNA. Wells containing only water and no DNA were also stained to obtain the background signal of the EvaGreen dye in the absence of DNA. Slides were covered to avoid exposure to light and incubated for 30 minutes at room temperature. The slides were then washed with water and dried under a flow of nitrogen gas.

### Slide imaging

Slides were fluorescently imaged using the InnScan1100 slide scanner (Innopsys). A 647nm laser was used to image the Sulfo-Cy5 fluorophore, a 532nm green laser was used to image the TAMRA fluorophore, and a 488nm laser was used to image the EvaGreen stain. DNA labelled samples with Cy5 or TAMRA fluorophores were imaged before EvaGreen staining in order to avoid co-excitation by the green laser.

### Data analysis

Images were analysed using ImageJ. The mean fluorescence intensity inside each well in both channels was extracted. The background signal was determined from the control replicates and subtracted from the TAMRA/Cy5 fluorescence signal in each sample well. To account for background noise in the EvaGreen signal (total DNA), a mean fluorescence signal of all wells containing EvaGreen and no DNA was calculated and subtracted from the EvaGreen signal in each sample well. The calculated epigenetic signal in each well was divided by the fluorescence intensity calculated in the EvaGreen channel of the same well in order to normalize the signal to the actual amount of DNA in the well. Next, the average and standard deviation for each sample were calculated over three to five replicates. In the case of 5hmC, the relative intensity was compared to the value of a calibration sample with a known 5hmC level, determined by LC-MS/MS. The final calculated value is the absolute 5hmC level in each sample.

## Data Availability

The raw data underlying this article are available upon request.

## Supplementary Data

Supplementary Data are provided.

## Acknowledgements

This work was supported by the European Research Council consolidator [grant number 817811] to Y.E;

Israel Science Foundation [grant number 771/21] to Y.E; The National Institute of health/ The National Human Genome Research Institute (NIH/NHGRI) [grant number R01HG009190] to Y.E; and Israel Innovation Authority and German Federal Ministry of Education and Research [NATI 61976 and 13GW0282B] to Y.E and E.W.

The authors declare no conflicts of interest

## Entry for the Table of Contents

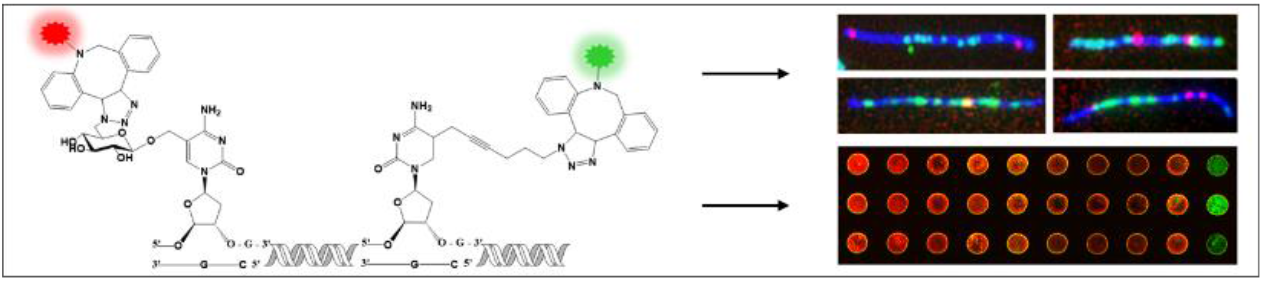

Simultaneous global labelling of 5-hydroxymethylcytosine and 5-methylcytosine by multi-colour fluorescent labelling. We apply a new methyltransferase enzyme specific for unmodified CpG to incorporate a modified cofactor that binds to a fluorophore by click chemistry. In combination with 5-hydroxymethylcytosine labelling *via* enzymatic glycosylation, we incorporate spectrally distinct colour for each epigenetic mark, enabling simultaneous quantification in different cancer types.

Institute and/or researcher Twitter usernames: @hagenom

## Notes

### Competing Interest Statement

The authors have declared no competing interest.

