## Supplementary Data for "Simultaneous global labeling (SiGL) of 5-methylcytosine and 5-hydroxymethylcytosine by DNA alkylation with a synthetic cofactor and engineered methyltransferase"

**Table of Contents**


1.1 Enzymatic characterization 3
1.1.1 Preparation of M.MpeI Q136A/N374A 3
1.1.2 Cofactor screening by modification/restriction assay 4
1.1.3 Quantification by RP-HPLC fragmentation assay 4
1.2 PCR amplification 7
1.2.1 Primer sequences for Lambda DNA amplicons 7
1.2.2 PCR procedure 8
1.3 Enzymatic labelling 8
1.3.1 Um- CpG labelling with MTaql 8
1.3.2 5hmC labelling 9
1.3.3 Control samples 9
1.4 Single-molecule imaging 9
1.4.1 Glass coverslips activation 10
1.4.2 Single-molecule imaging with an epi-fluorescence microscope 10
1.5 Biological samples 11
1.5.1 Colorectal cancer (CRC) samples 11
1.5.2 Hematologic cancer samples 11
2.1 Technical validation 12
2.2 Limit of detection 13
2.3 Chemoenzymatic labelling of unmodified CpGs 14
2.4 Labelling efficiency 15
4 MTaql labelling of um-CpGs in colon samples 16
5 Fluorescent signal intensity in colon samples 16
5.1 5hmC in colon 17
5.2 um-CpG in colon 17
6.1 5hmC in hematological samples 18
6.2 um-CpG in hematological samples 19
References 20

### 1 Experimental procedures

#### 1.1 Enzymatic characterization

HPLC-purified oligodeoxynucleotides (ODN) were purchased from IDT. Proteinase K was obtained from Qiagen, DBCO-PEG4-TAMRA and UDP-6-azidoglucose were from Jena Bioscience, calf intestinal alkaline phosphatase (CIAP) was from Takara Bio and nuclease P1, as well as T4 phage β-glucosyltransferase (T4-BGT), were from NEB. Poros HS 50 column material was from GE-Healthcare, Ni-NTA-agarose column material from Qiagen, and Prontosil C-18 HPLC column from Bischhoff.

##### 1.1.1 Preparation of M.MpeI Q136A/N374A

*Protein expression:* Site-directed mutagenesis was used to introduce point mutations Q136A and N374A into M.MpeI gene within the pET28a-M.MpeI expression plasmid[1]. A colony of *E. coli* K12 SG1709 cells (Scarab Genomics LLC) containing pET28a-M.MpeI Q136A/N374A plasmid was cultured in LB medium (0.8 mL) containing kanamycin (35 mg/L) by incubation at 37 °C and 250 rpm for 6 h. An overnight culture (50 mL, LB medium with 35 mg/L kanamycin) was inoculated with the starting culture (0.8 mL) and incubated by shaking at 250 rpm and 37 °C overnight. LB medium (1 L) with kanamycin (35 mg/L) was inoculated with the overnight culture (25 mL) and incubated at 37 °C and 200 rpm until an optical density $OD^{600}$ of about 0.6 was reached. IPTG (0.4 mL, 1 M) was added to the culture and incubation continued at 37 °C and 200 rpm for 20 h. Cells were harvested by centrifugation at 3,500 rpm and 4° C for 15 min. The supernatant was decanted and the cell pellet was stored at -20 °C.
*Protein purification:* Cells were resuspended in lysis buffer (50 mL, 50 mM Tris-HCl, 300 mM NaCl, 5% glycerol, 1 mM DTT, pH 7.5) and disrupted by sonication on ice for 4 × 1 min (output control 8, duty cycle 60 %) with breaks of 1 min. The suspension was mixed with Triton X-100 (2.4 mL, 25%), incubated on ice for 10 min, and centrifuged (16,500 g, 4 °C, 1 h). Cleared lysate was obtained after decantation, diluted with an equal volume of buffer (50 mM Tris-HCl, 500 mM NaCl, 1 mM imidazole, pH 7.5), and applied to a Ni-NTA-agarose column (2 mL) which was equilibrated with the same buffer (20 mL). The column was treated with wash buffer (20 mL, 50 mM Tris-HCl, 500 mM NaCl, 10 mM imidazole, pH 7.5). Proteins were eluted with a step gradient of imidazole (2 mL 50 mM, 2 mL 100 mM, 2 mL 150 mM and 16 mL 250 mM) in buffer (50 mM Tris-HCl, 500 mM NaCl, pH 7.5). Fractions (2 mL) were collected and analyzed by SDS-PAGE. Fractions containing M.MpeI Q136A/N374A were combined and diluted with low-salt buffer (50 mM Tris-HCl, 100 mM NaCl, pH 7.5). Proteins were loaded on a cation-exchange column (20 mL, Poros HS 50) which was equilibrated before with cofactor releasing buffer (100 mL, 6.7 mM MES, 6.7 mM NaOAc, 6.7 mM HEPES, 200 mM KCl, 0.2 mM DTT, 10% glycerol, pH 6.0). The column was washed with cofactor-releasing buffer (6 L) in order to wash away bound natural cofactor from the protein. M.MpeI Q136A/N374A was eluted with a linear salt gradient from 100 mM to 1000 mM NaCl. Fractions containing high protein concentration (650–1000 mM NaCl) were combined, placed in a dialysis tubing (molecular weight cut-off 12,000–14,000), and dialyzed at 4 °C in dialysis buffer (2 L, 50 mM Tris-HCl, 500 mM NaCl, 10 mM 2-mercaptoethanol, 50% glycerol, pH

7.5) for 18 h. 30 mg of purified M.MpeI Q136A/N374A were obtained from 1 L cell culture and stored at -20° C.

##### 1.1.2 Cofactor screening by modification/restriction assay

Solutions (20 µL) containing a derivative of Litmus28*i* plasmid (1 µg), M.MpeI WT or M.MpeI Q136A/N374A (5 µM, 1 eq. regarding CpG sites) and AdoMet (80 µM) or AdoMet analogue (80 µM for epimeric pure compounds and 160 µM for epimeric mixtures at sulfur) in reaction buffer (50 mM Tris-HCl, 100 mM NaCl, 10 mM 2-mercaptoethanol, 0.1 mg/mL BSA, 5% glycerol, pH 7.5) were incubated at 37 °C for 1 h. Proteinase K (600 µAU) was added and incubation continued at 37 °C for 1 h. Plasmid DNA of each sample was purified with QIAquick® PCR purification kit (Qiagen), eluted in EB buffer (20 µL), supplemented with R.BstUI (10 U) in CutSmart buffer (30 µL, 50 mM Tris-OAc, 10 mM $Mg(OAc)_2$, 50 mM KOAc, 0.1 mg/mL BSA, pH 7.9) and incubated at 60 °C for 1 h. The reaction mixtures were treated with proteinase K (600 µAU), incubated at 37 °C for 1 h and 6× loading buffer (10 µL, 0.25% bromphenol blue, 30% glycerol) was added. Samples (10 µL) were analyzed by agarose gel (1%) electrophoresis in TBE buffer (0.5×) containing GelRed (0.05 µL stock solution/mL gel). DNA bands were visualized with a UV transilluminator (312 nm) and documented with a CCD camera equipped with a filter (540 ± 50 nm).

##### 1.1.3 Quantification by RP-HPLC fragmentation assay

###### *1.1.3.1 Labelling cytosines in CpG sites*

*Transfer of azide groups (1. step):* A solution (200 µL) of M.MpeI Q136A/N374A (115 ng/µL, 2.5 µM) in reaction buffer (50 mM Tris-HCl, 100 mM NaCl, 10 mM 2-mercaptoethanol, 0.1 mg/mL BSA, 5% glycerol, pH 7.5) containing duplex ODN (hybridization of 5´-ATT ATT ATT ATT AG**C G**CA TTA TTA-3´ and 5’-TAA TAA TG**C G**CT AAT AAT AAT AAT-3’, 2.5 µM CpG sites) and AdoYnAzide[2] (80 µM) was prepared. To exclude oxygen deionized water was degassed before by filtration through a regenerated cellulose filter (0.2 µm) and the buffer (10× without 2-mercaptoethanol) was degassed by repeated (three times) freezing using liquid nitrogen, applying static vacuum, and thawing (freeze-pump-thaw degassing). 2-Mercaptoethanol was added afterward. The reaction mixture was incubated at 37 °C overnight. Proteinase K (1 µL, 20 µg/µL) was added and incubation continued at 37 °C for 1 h, followed by heat inactivation at 80 °C for 30 min.
*Fluorescence click labelling (2. step):* Half of the reaction mixture (100 µL) was supplemented with DBCO-PEG4-5/6-TAMRA in DMSO (1.6 µL, 10 mM, mixture of 5- and 6-isomer) to give a final concentration of 160 µM. The click reaction was performed at 37 °C overnight.
*Enzymatic fragmentation:* Reaction mixtures were purified using NAP-5 columns following the instructions of the manufacturer. Nuclease P1 buffer (20 mM NaOAc, 0.2 mM $ZnSO_4$, pH 5.3) was used for column equilibration (10 mL) and elution (500 µL). Purified duplex ODN were supplemented with nuclease P1 (3 µL, 10 U/µL) and incubated at 37 °C for 2 h. CIAP (1.5 µL, 0.6 U/µL), 10× CIAP buffer (60 µL, 500 mM Tris-HCl, 10 mM $MgCl_2$, pH 9.0), and deionized water were added to a total volume of 600 µL and samples incubated at 37 °C overnight.

*HPLC analysis:* Nucleosides were separated by reverse-phase (RP) HPLC (Prontosil C-18 AQ column, 250 × 4.6 mm, 5 µm, 120 Å, equipped with a C-18 pre-column, 8.0 mm × 4.0 mm, 5 µm, 120 Å) using an acetonitrile gradient (3.5–5.7% in 20 min and 5.7–70% in 22 min) in aqueous TEAAc (10 mM) with a flow of 1 mL/min and detection at 260 nm/272 nm or 260 nm/555 nm for runs with TAMRA-labeled nucleosides.
*Quantification of nucleosides:* Peaks of nucleosides at 260 nm were integrated using Empower 2 software (Waters) and areas were normalized by dividing them by the respective extinction coefficients at 260 nm (dC: 7,009 mol $L^{-1}$ $cm^{-1}$; dG: 11,715 mol $L^{-1}$ $cm^{-1}$; dT: 8,902 mol $L^{-1}$ $cm^{-1}$; dA: 15,663 mol $L^{-1}$ $cm^{-1}$; 5-azido-dC = 5mdC: 5,435 mol $L^{-1}$ $cm^{-1}$). Normalized areas of dT were divided by the number of dT residues in the duplex ODN to obtain a normalized area for one nucleoside. Normalized areas for the other nucleosides were divided by this value to give experimental amounts for all nucleosides. The yield for 5-azido-dC was calculated by dividing the amount of 5-azido-dC by the sum of dC and 5-azido-dC multiplied by 2 (four dC/two target dC). The yield for 5-TAMRA-dC was calculated from the decrease of 5-azido-dC.

###### *1.1.3.2 Sequence specificity of M.MpeI Q136A/N374A*

Solutions of M.MpeI Q136A/N374A (230 ng/µl, 5.0 µM) in reaction buffer (50 mM Tris-HCl, 100 mM NaCl, 10 mM 2-mercaptoethanol, 0.1 mg/mL BSA, 5% glycerol, pH 7.5) containing either a duplex ODN with CpG site (hybridization of 5´-GTC AG**C** **G**CA TCC -3´ and 5'-GGA TG**C** **G**CT GAC-3', 5 µM, 5 µM CpG sites) or without CpG site (hybridization of 5´- GTC AGG CCA TCC -3´ and 5'-GGA TGG CCT GAC-3', 5 µM) and AdoMet (80 µM) were prepared. Oxygen was excluded as described above (4.1) and the reaction mixtures were incubated at 37 °C for 1 h. Proteinase K (1 µL, 20 µg/µL) was added to the reaction mixtures and incubation continued at 37 °C for 1 h, followed by heat inactivation at 80 °C for 30 min. Enzymatic fragmentation, HPLC analysis, and quantification of nucleosides followed the protocol given above (1.1.3.1) except that yields for 5mC were calculated by dividing the amount of 5mC by the sum of dC and 5mC multiplied by 4 (eight dC/two target dC for the specific duplex).

###### *1.1.3.3 Labelling 5-hydroxymethylcytosine*

A solution (100 µL) containing duplex ODN with 5hmC (20 µM, hybridization of 5´-TGT CAG 5hmCGC ATG A-3´ and 5´-TCA TGC GCT GAC A-3'), UDP-6-azidoglucose (40 µM) and T4 phage β-glucosyltransferase (T4-BGT) (1.72 U/µL) in CutSmart buffer were incubated at 37 °C overnight (1. step). 50 µL of this solution were supplemented with DBCO-PEG4-TAMRA (160 µM) and incubation was continued at 37 °C overnight (2. step). Both samples (50 µL) were treated with deionized water (50 µL) as well as nuclease P1 buffer (400 µL) and purified using NAP-5 columns. Nuclease P1 buffer was used for column equilibration (10 mL) and elution (500 µL). Both samples were heated to 100 °C for 10 min and placed on ice for 2 min before nuclease P1 (1 µL, 1 U/µL) was added. After incubation at 37 °C for 2 h 10× CIAP buffer (60 µL), CIAP (3 µL, 0,16 U/µL), and deionized water were added to total volumes of 600 µL and samples were incubated at 37 °C overnight. Nucleosides were separated by RP-HPLC and quantified as described above (1.1.3.1).

**Table S1.** AdoMet and analogues for cofactor screening with M.MpeI WT and M.MpeI Q136A/N374A.

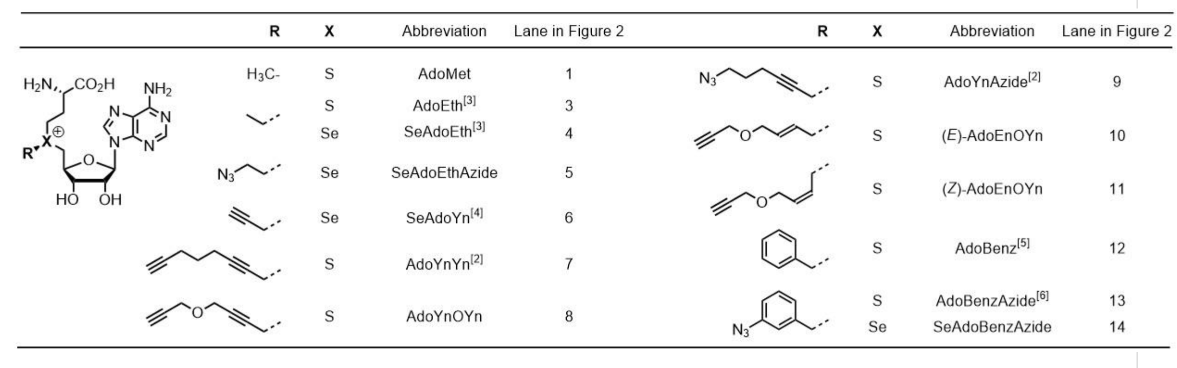

| R | X | Abbreviation | Lane in Figure 2 | R | X | Abbreviation | Lane in Figure 2 |
|---|---|---|---|---|---|---|---|
| $H_3C$- | S | AdoMet | 1 | | S | AdoYnAzide[2] | 9 |
| | S | AdoEth[3] | 3 | | S | (*E*)-AdoEnOYn | 10 |
| | Se | SeAdoEth[3] | 4 | | S | (*Z*)-AdoEnOYn | 11 |
| | Se | SeAdoEthAzide | 5 | | S | AdoBenz[5] | 12 |
| | Se | SeAdoYn[4] | 6 | | S | AdoBenzAzide[6] | 13 |
| | S | AdoYnYn[2] | 7 | | Se | SeAdoBenzAzide | 14 |
| | S | AdoYnOYn | 8 | | | | |

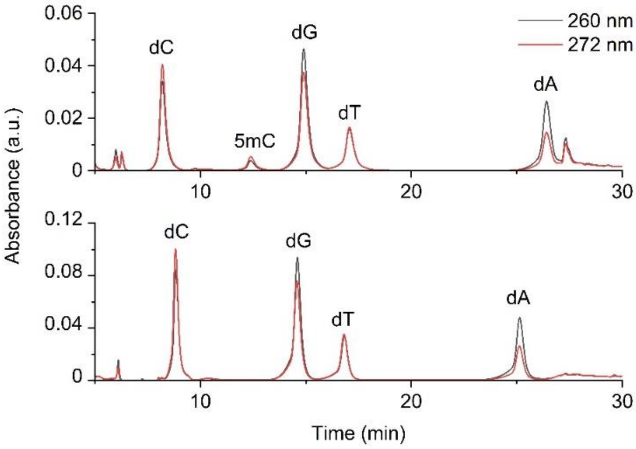


**Figure S1.** RP-HPLC analysis of nucleosides obtained from incubations of M.MpeI Q136A/N374A with a specific duplex DNA containing one CpG site (upper) or a non-specific duplex ODN without CpG site (lower) followed by enzymatic fragmentations.

**Table S2.** Quantification of cytosine methylation with M.MpeI Q136A/N374A by RP-HPLC analysis (see Figure S1). Amounts of nucleosides obtained after incubations with a specific duplex DNA containing one CpG site or a non-specific duplex ODN without CpG site.

| Duplex ODN | dC | dG | dT | dA | 5mdC | Yield [%] |
|---|---|---|---|---|---|---|
| One CpG | 8.34 | 8.97 | 4.00 | 3.73 | 0.97 | 42 |
| No CpG | 8.93 | 8.48 | 4.00 | 3.77 | 0 | 0 |

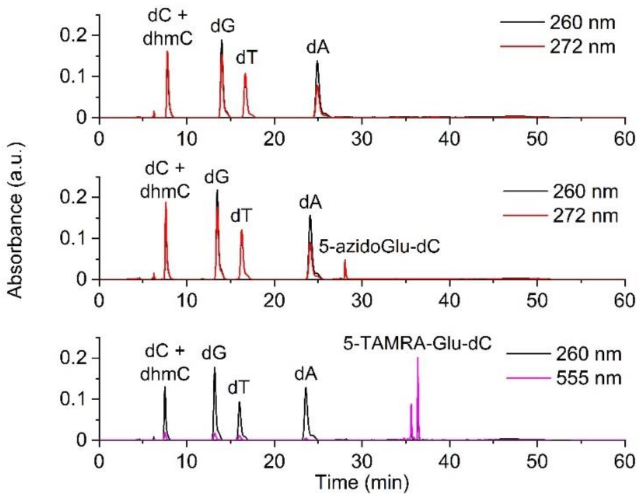


**Figure S2.** 5hmC labelling analysis by RP-HPLC of nucleosides obtained after enzymatic fragmentation of duplex ODN. Top: Control reaction with duplex ODN containing one 5hmC residue. Middle: Modification of the duplex ODN with T4-BGT and UDP-6-azidoglucose (1. step). Bottom: Click reaction of azido-modified duplex ODN with DBCO-PEG4-TAMRA (2. step). Two signals are observed for 5-TAMRA-Glu-dC because the strain-promoted azide-alkyne cycloaddition (SPAAC) can give two regioisomers.

**Table S3.** Quantification of 5hmC labelling analysis by RP-HPLC analysis (see Figure S2). Amounts of nucleosides obtained after duplex modification with T4-BGT and UDP-6-azidoglucose (1. step) followed by click labelling with DBCO-PEG4-TAMRA (2. step).

| | dC + 5hmdC | dG | dT | dA | 5-azido-Glu-dC | 5-TAMRA-Glu-dC | Yield [%] |
|---|---|---|---|---|---|---|---|
| Control | 7.0 | 7.0 | 6.0 | 6.0 | - | - | |
| 1. step | 6.0 | 7.0 | 6.3 | 5.8 | 1.1 | - | 100 |
| 2. step | 6.1 | 7.0 | 6.4 | 6.0 | 0.1 | 1.0 | 91 |

#### 1.2 PCR amplification

##### 1.2.1 Primer sequences for Lambda DNA amplicons

**Table S4.** Primer sequences for Lambda DNA amplicons

| Amplicon size (bp) | Forward (5'→3') | Reverse (3'→5') |
|---|---|---|
| 222+16CpG sites | GTGATGCCGAGAACTTTATG | AAGCAGCAAGTTCATCATAAC |

| | | |
|---|---|---|
| 204+2 fluorophores | Cy5/TAMRA-GTGCAATGAAGCCAAGTTAGAA | TAMRA-TCTTTTGTAATAGTGTCTTTTGTGT |
| 408+1 fluorophore | Cy5/TAMRA-GTGCAATGAAGCCAAGTTAGAA | GCCCACCCAGCAAAATTC |
| 816 + 1 fluorophore | Cy5/TAMRA-GTGCAATGAAGCCAAGTTAGAA | ACGGTTAGCCAGGCTCG |
| 1632 + 1 fluorophore | Cy5/TAMRA-GTGCAATGAAGCCAAGTTAGAA | GTCATCTTTTAACTCCATATACCGC |
| 202 + 1 CpG site | GATTTGGCCATACTACTAAATCC | GGAAATGTTAAAATTGGTTTTGTAG |
| 204 + 2 CpG sites | AGAATTTTTAGCCCAAGCCA | CACTTGCTCAAATGCTGCAT |
| 204 + 4 CpG sites | GTGCAATGAAGCCAAGTTAGAA | TCTTTTGTAATAGTGTCTTTTGTGT |
| 210+8CpG sites | GTGTATTACCGGTTTGCTAC | AGTTTTTTCATGACTTCCCTC |
| 222+16CpG sites | GTGATGCCGAGAACTTTATG | AAGCAGCAAGTTCATCATAAC |

##### 1.2.2 PCR procedure

PCR was used to synthesize 200-1600 base pairs (bp) long DNA fragments using lambda DNA as a template. Some of the primers contained one fluorophore for assay characterization and validation. For PCR amplification 100 ng of lambda DNA (New England biolabs) was mixed with 0.5µL of forward and reverse primers, 12.5 µL of HS Taq mix red (PCRBIO), and ultrapure water to a final volume of 25 µL. The PCR program was as follows; initialization at $94^0$C for 2 minutes and then 30 cycles of denaturation at $95^0$C for 20 seconds, annealing at $53^0$C for 20 seconds, and elongation at $72^0$C for 30 seconds to achieve 200-400bp fragments and 90 seconds for 800-1600bp DNA fragments. After the 30 cycles, a final elongation was performed at $65^0$C for 2 minutes. After PCR amplification the samples were purified using QIAquick PCR Purification Kit (Qiagen).

#### 1.3 Enzymatic labelling

##### 1.3.1 Um- CpG labelling with MTaql

Um-CpGs in the context of TCGA sequence were fluorescently labelled via a two-step chemoenzymatic reaction. In each reaction tube, 500 ng of DNA were mixed with 2.5 µL 10X buffer 4 (New England Biolabs), AdoYnN3 to a final concentration of 40 µM, MTaql (NEB) to a final concentration of 0.0078 mg/mL, and ultrapure water to a final volume of 25 µL. The reaction mixture was incubated for 1 hour at 60 °C. Then, 20 µg of Proteinase K (PK) (Sigma) was added and incubated for 2 hours at 45 °C. Following inactivation, Dibenzocyclooctyl (DBCO)-PEG4-5/6- TAMRA or Dibenzocyclooctyl (DBCO)-Sulfo-Cy5 (Jena Bioscience, Jena, Germany) was added to a final concentration of 250 µM and then incubated overnight at $37^0$C. The labelled DNA samples were purified from excess fluorophores using Oligo Clean & Concentrator columns (Zymo Research), according to the manufacturer's recommendations, with three washing steps and two elution steps for optimal results.

##### 1.3.2 5hmC labelling

5hmC residues were fluorescently labelled via a two-step chemoenzymatic reaction[7-9]. In each reaction tube, 1 µg of genomic DNA was mixed with 3 µl of 10X buffer 4 (New England Biolabs), uridine diphosphate-6-azide-glucose (UDP-6-N3- Glu) to a final concentration of 45 µM, 2 µl (20 units) of T4 phage β[1]glucosyltransferase (T4-BGT, New England Biolabs), and ultrapure water to a final volume of 30 µL. The reaction mixture was incubated overnight at 37 °C. The following day, Dibenzocyclooctyl (DBCO)-PEG4-5/6-TAMRA Dibenzocyclooctyl (DBCO)-Sulfo-Cy5 (Jena Bioscience, Jena, Germany) was added to a final concentration of 150 µM, and the reaction was incubated overnight at 37 °C. The labeled DNA samples were purified from excess fluorophores using Oligo Clean & Concentrator columns (Zymo Research), according to the manufacturer's recommendations, with three washing steps and two elutions for optimal results. For best yield, not more than two micrograms of DNA (two identical reaction tubes combined) were loaded on one column. Samples were kept at 4 °C until analyzed.

##### 1.3.3 Control samples

We performed an identical reaction without the enzyme (M.MpeI (dm), MTaql, or T4-βGT, depending on the desired labelling) for control samples. The result is a non-labelled DNA sample with fluorescence signals originating from remaining free fluorescent dye molecules. This residual signal, representing the assay's noise level, was subtracted during analysis to represent the signal derived from labelled 5-hmC residues reliably.

#### 1.4 Single-molecule imaging

##### 1.4.1 Glass coverslips activation

Single-molecule imaging was performed using a microfluidic platform that enables capturing and stretching of fluorescently labeled DNA molecules on a glass surface. Because DNA is negatively charged (due to its phosphate backbone), it is possible to attach it to a glass surface by coating it with a positively charged substance, creating an electrostatic binding. In this work, glass coverslips were activated according to the following protocol. 22X22 glass coverslips surface were cleaned and enriched for hydroxyl groups by immersing them in a 300mL mixture of nitric acid and hydrochloric acid at a ratio of 2:1 respectively and incubated overnight. After incubation, the coverslips were thoroughly washed twice with DDW and twice with 96% ethanol and then dried under a nitrogen gas flow. The dried coverslips were immediately immersed in 300mL aqueous solution containing two types of silanes; 750µL of N-trimethoxysilylpropyl-N,N,N-trimethylammonium chloride (Gelest) to a final concentration of 8.99µM and 250 µL of vinyltrimethoxysilane (Gelest) to a final concentration of 5.45µM. Then the coverslips were incubated at $65^0$C for 1050 minutes. After incubation, the coverslips were thoroughly washed twice with DDW and twice with 96% ethanol and stored in 96% ethanol at $4^0$C. Right before using, the coverslip is rinsed with DDW and ethanol and dried under a flow of nitrogen gas. The coverslip is then placed on a glass microscope slide (Color Frost, Bar Naor). DNA sample deposition for imaging was done by loading the sample on the contact area of the slide and the coverslip. DNA is stretched by capillary tension that pulls the DNA solution and binding force between the negatively charged DNA and the positively charged moieties of the amino-silanes on the coverslip. During the flow, one of the ends of the DNA binds to the hydrophobic vinyltrimethoxysilane. This occurs due to spontaneous denaturation of the dsDNA ends, which exposes the hydrophobic nitrogenous bases. The rest of the strand is pulled by the capillary flow and is anchored by the positively charged amino silanes until the DNA molecule is bound to the surface.

##### 1.4.2 Single-molecule imaging with an epi-fluorescence microscope

After loading the DNA sample, the glass microscope slide was placed in an epifluorescence microscope (TILL Photonics) with the coverslip facing the objective. The objective used in this work, is a 100x oil objective (UPlanSApo, ∞/0.17/FN26.5/ N.A 1.4, Olympus) and an EMCCD camera (iXon888, Andor) for imaging. The autofocus feature used to image multiple fields of view (FOVs) is based on infrared reflection from the sample's surface. Sequential images were taken in each FOV to visualize the labeled DNA's backbone and the epigenetic modifications along its contour. The fluorophores and filters used for imaging were YOYO-1 ex485/20, and em525/30 for DNA. ATTO647N ex640/4, em684/24 and ATTO550 ex563/9, em578/16 for um-CPG and 5hmC. In this work, single-molecule imaging was used to verify and quantify the simultaneous labelling of 5hmC and 5mC on the same DNA sample in the SiGL experiments. Sample preparation and staining of the DNA backbone were done by mixing labeled DNA to a final concentration of 0.2 ng/µL, 12.5 µL of 2M DDT, 0.23 µL of 20 µM YOYO-1 intercalator dye, and 1X TE buffer to a final volume of 50 µL. From this solution, 7 µL was taken for fluorescent imaging.

#### 1.5 Biological samples

##### 1.5.1 Colorectal cancer (CRC) samples

DNA samples from CRC tumors and tumor-adjacent tissues and DNA from healthy individuals were purchased for analysis (OriGene, DYN diagnostics).

##### 1.5.2 Hematologic cancer samples

Whole peripheral blood samples from patients with Chronic lymphocytic leukaemia and healthy individuals were received as a part of a collaboration with Dr. Maya Koren-Michowitz and Dr. Roni Golan-Lavi from Shamir medical center (Assaf HaRofe). The samples were collected with informed consent for research use and approved by the Tel-Aviv University and Shamir Medical Center ethical review boards, in accordance with the declaration of Helsinki.

The steps of the isolation of peripheral blood mononuclear cells (PBMCs) are as follows:

0.5mL of peripheral blood were frozen in 1mL cryogenic freezing tubes with a screw cap at -80C for later DNA extraction. The rest of the samples were separated for PBMCs. Each Blood was diluted with one volume of PBSX1 and then Ficoll-Paque (GE Healthcare) was added to a 50mL conical tube in a 3:4 ratio to the blood samples and then the diluted blood was layered slowly and carefully on top of the Ficoll-Paque. The samples were then centrifuged at 400Xg for 30 minutes at 200C; acceleration 6, deceleration 1. After centrifuge, the plasma layer was removed without disturbing the PBMC layer and the interface layer containing the PBMCs was transferred into a new 50mL conical tube. The cells were washed with 3X original blood volume of PBSX1 by pipetting up/down using a serological pipet and then centrifuged at 200Xg for 10 minutes at 200C and the supernatant was discarded. The cells were re-suspended in 10mL PBSX1 and mixed thoroughly. To count the cells, 10µL of the sample was added to the hemocytometer and placed in a chamber in a microscope under a 10X objective. The cells were counted in the large, central gridded square (4x4 squares, 1mm2) and multiple by 104 to estimate the number of cells per milliliter. The final number of cells was calculated as follows:

$$\textit{Total Number of cells} \\ = \textit{Number of cells in } 4X4 \textit{ square X Dilution factor X } 10^4 \textit{ X Total sample Volium } (mL)$$

**Equation S1:** Total number of cells calculation.

Then the re-suspended cells were centrifuged at 200Xg for 10 minutes at 200C and the supernatant was discarded. Cells were re-suspended in 1mL freezing buffer (90% FBS with 10% DMSO). For each sample two cryogenic freezing tubes with a screw cap were prepared; one for freezing 4 million cells and one for the rest. After transferring 4x106 cells to one tube and the rest to the second tube, each tube was filled with freezing buffer to a final volume of 1mL and frozen at -80C for later DNA extraction (omega, BIO-TEK).

### 2 Assay validation

#### 2.1 Technical validation

To design a multi-sample array platform for rapid quantification of 5mC and 5hmC, we start by characterization and validating the system. First, to ensure that the slide assay is reliable across different DNA lengths, we prepared a set of DNA samples with one fluorophore and varying lengths (table S4). We then applied equal amounts of DNA from each sample on the activated multi-sample array slide and measured the fluorescence intensity values. These values were normalized to the sample with the highest intensity and then to the expected theoretical values based on the length of the DNA fragments (equation S2).

$$\%\textit{labeling } (\textit{theoretical}) = \left(\frac{\textit{Number of fluorophores}}{\textit{Number of total nucleobases}}\right) X\ 100\%$$

**Equation S2.** Theoretically expected labelling percentage.

This procedure was performed using two different fluorescent dyes, TAMRA and Cy5, to ensure the method is insensitive to the dye used. The results from both fluorophores in this study corroborate the expected relative fluorescence values, and the results are consistent as shown by the small error bars in Figure S3.

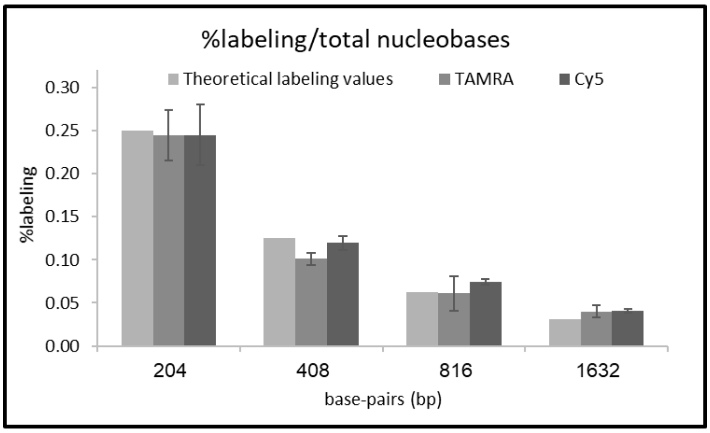


**Figure S3.** Assay characterization and validation. Normalized fluorescent intensity of the samples. The error bars stand for the standard deviation of five replicates.

#### 2.2 Limit of detection

To test the assay's limit of detection (LOD) and calculate the minimal amount of sample (analyte) that can be reliably detected by the slide assay we measured a decreasing amount of labelled DNA to produce a LOD curve that reports the sensitivity in units of the percent of labelled nucleotides out of the total number of nucleotides. To achieve this, we used a 400bp PCR product amplified either with standard primers or with a forward primer containing one TAMRA fluorophore (table S4). Then we prepared 12 different dilutions for labelling percentages ranging from 0.124% (400bp with one fluorophore) to 0% (400bp with no fluorophore) with equal DNA concentration in each sample by mixing the two PCR products in different ratios. To account for background noise we used a control sample composed of DNA that received the labelling reaction without the enzyme. The background signal from this sample was added to each of the synthetic samples used in the LOD curve. A standard curve of 12 points was created with the relative intensity values obtained from the slide scanner measurements (response) and the expected labelling percentage based on the serial dilutions (Figure S4). We used the calibration curve method based on the standard deviation of the response (σ) and slope (s) of the calibration curve, making it more accurate. The Standard deviation can be calculated for the Y-intercept and used to calculate the LOD for a standard curve using a regression function[5]. We calculated a LOD of 0.0024%, which is lower than the 5hmC content in blood, the tissue with the lowest levels of 5hmC among the different tissues in the human body. Moreover, the LOD value is even lower than that of the blood cancer patients, highlighting the assay's sensitivity and ability to reliably measure and distinguish between healthy and sick individuals even at these low 5hmC levels.

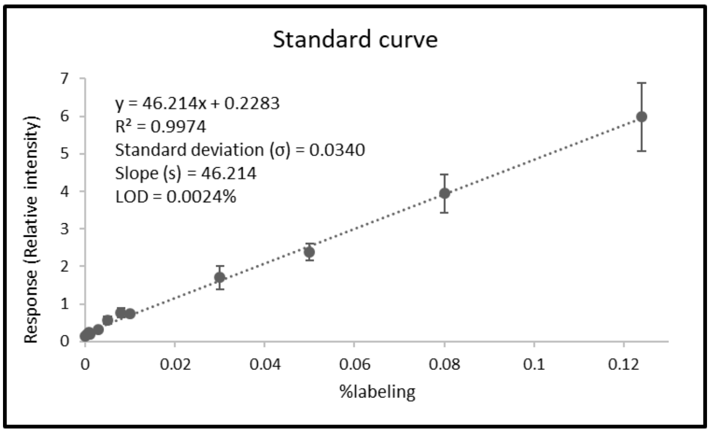


**Figure S4.** Standard curve created with 400bp PCR products containing a known labelling percentage. The graph shows relative intensity measurements obtained from the slide scanner versus the expected labelling percentage based on serial dilutions.

#### 2.3 Chemoenzymatic labelling of unmodified CpGs

To verify that the assay fluorescence readout is linear with respect to the number of available CpGs we used a PCR amplicon with varying amount of CpGs (table S4). The amplification was performed using specific forward and reverse primers to achieve DNA fragments of the same length (~200bp) with a different number of CpG sites in each one (1,2,4,8 and 16 CpGs). We performed the M.MpeI (dm) labelling reaction for um-CpGs and applied equal amounts of DNA to the activated slides. The labelling reaction was performed using both DBCO-TAMRA and DBCO-Cy5 to verify that there is no difference between the fluorophores' attachments to the slide. The results show a linearly increasing signal with the increased number of CpGs for both fluorophores for the M.MpeI (dm) labelling reaction (Figure S5).

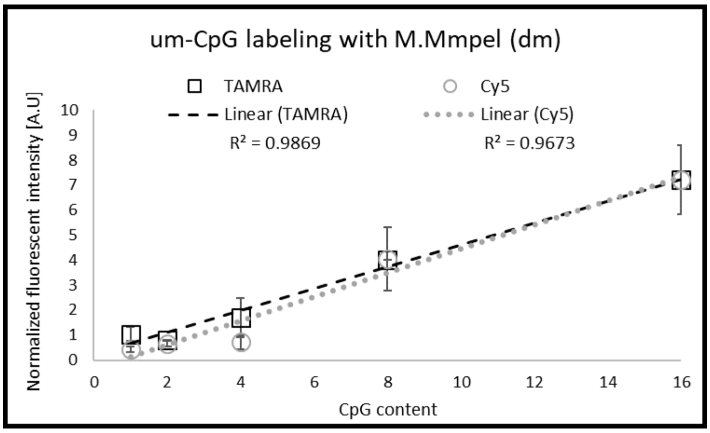


**Figure S5.** Normalized fluorescent intensity of the samples to the total amount of DNA and the theoretical labelling percentage. The error bars stand for the standard deviation of five replicates. 200bp products with a different number of CpG sites in each one (1,2,4,8 and 16 CpGs) labelled with M.MpeI (dm) and either DBCO-TAMRA or DBCO-Cy5 fluorophore.

#### 2.4 Labelling efficiency

CG is a palindromic sequence generating two methylation sites on opposite strands. In principle, M.MpeI (dm) incorporates the modified cofactor on both strands of the DNA. To test the labelling efficiency, we prepared a 200bp PCR product with one CpG site on each DNA strand (table S4). The PCR product was then labelled with M.MpeI (dm) and compared to a PCR product amplified using two fluorescent primes (forward and reverse), containing one a TAMRA fluorophore on each one. This PCR product represents 100% labelling efficiency (equivalent to both CpGs fluorescently labeled). In the case of DBCO-Cy5, the labeled sample was compared to a PCR product amplified using only one fluorescent primer containing one Cy5 fluorophore, and this represented 50% labelling. The results point to a labelling efficiency of 15% and 12% for the TAMRA and Cy5 fluorophores, respectively (Figure S6). Although this labelling efficiency is lower than the desired efficiency, it is sufficient for reliable results using the slide assay for global detection and quantification of um-CpG.

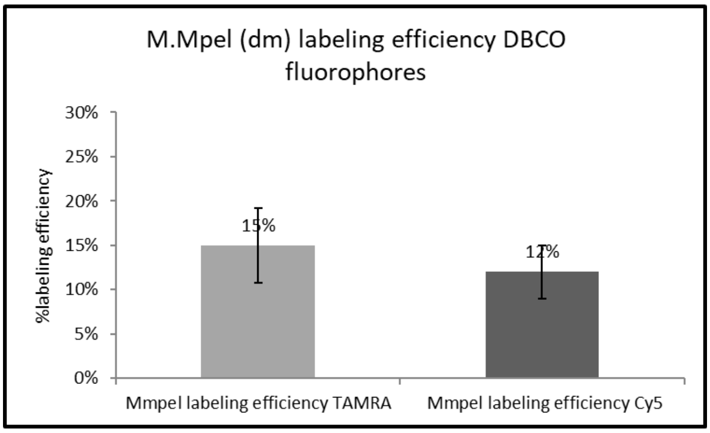


**Figure S6.** 200bp DNA fragment with one CpG site labelled with M.MpeI (dm) enzyme compared to a 200bp PCR product amplified using one Cy5 fluorophore, representing 50% labelling efficiency (right). 200bp DNA fragment with one CpG site labeled with M.MpeI (dm) enzyme and TAMRA fluorophore compared to a 200bp PCR product amplified using two TAMRA fluorophores, representing 100% labelling efficiency (left). The error bars stand for the standard deviation of three to five replicates.

### 3 Comparison between Mmpel (dm) Vs. MTaql methyltransferase

To compare the M.MpeI (dm) labelling to that of the previously published MTaql enzyme[11,12] we labelled unmethylated lambda DNA for um-CpGs using each one of the methyltransferase enzymes and then applied the labelled samples onto the activated slide. The labelling reaction was performed using DBCO-TAMRA fluorophore. To verify the results, we experimented twice with comparable results (5.56±1.37 in the first experiment and 6.39±2.26 in the second experiment). The results presented in figure S7 represent the average fluorescence intensity of the two experiments.

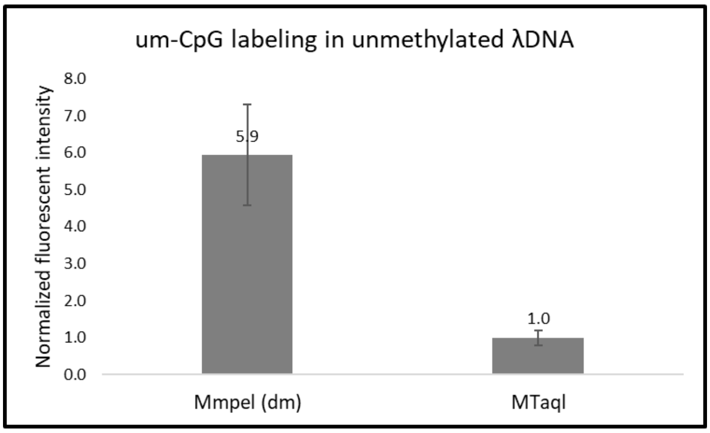


**Figure S7**. Normalized fluorescence intensity of um-CpG labelling with M.MpeI (dm) (left) and MTaql (right). The error bars stand for the standard deviation of two experiments, each one containing three to five replicates.

#### 4 MTaql labelling of um-CpGs in colon samples

To further validate the methylation results in the colon, we repeated the um-CpG labelling using the MTaql enzyme. This experiment aims to verify that both enzymes produced similar trends. The two highest intensity samples (colon cancer 2) were normalized to 1 in order to bring the two data sets to the same intensity scale. The plot in Figure S8 shows that M.MpeI (dm) labelling follows the MTaql labelling trend.

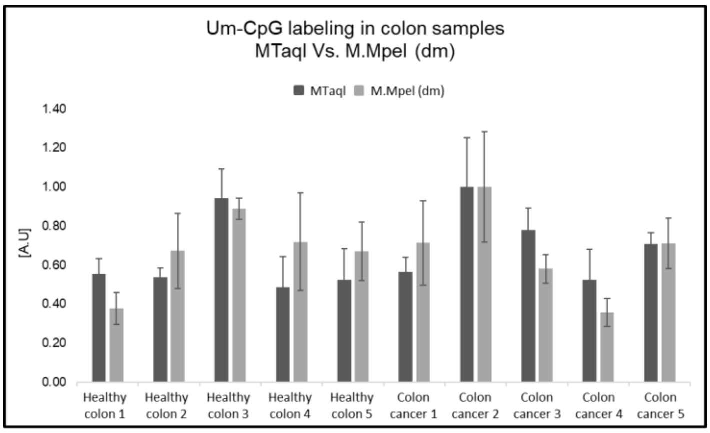


**Figure S8.** MTaql and M.MpeI (dm) labelling in colon. Normalized fluorescence intensity of healthy colon (N=5) and colon cancer (N=5) labeled with M.MpeI (dm) (light gray) and MTaql (dark gray). The error bars stand for the standard deviation of three to five replicates.

#### 5 Fluorescent signal intensity in colon samples

#### 5.1 5hmC in colon

**Table S5.** A table containing the normalized fluorescent signal intensity (epigenetic intensity/DNA intensity obtained from the slide scanner)

| Healthy colon (N=18) | | Colon tumor (N=20) | |
|---|---|---|---|
| Average of three to five replicates | Standard deviation | Average of three to five replicates | Standard deviation |
| 0.184 | 0.068 | 0.094 | 0.014 |
| 0.379 | 0.107 | 0.075 | 0.012 |
| 0.216 | 0.087 | 0.070 | 0.008 |
| 0.171 | 0.060 | 0.087 | 0.014 |
| 0.293 | 0.089 | 0.033 | 0.009 |
| 0.099 | 0.049 | 0.076 | 0.006 |
| 0.260 | 0.098 | 0.102 | 0.025 |
| 0.308 | 0.122 | 0.076 | 0.003 |
| 0.298 | 0.103 | 0.106 | 0.012 |
| 0.515 | 0.103 | 0.093 | 0.004 |
| 0.674 | 0.326 | 0.276 | 0.044 |
| 0.439 | 0.186 | 0.186 | 0.025 |
| 0.347 | 0.116 | 0.178 | 0.043 |
| 0.305 | 0.032 | 0.063 | 0.012 |
| 0.284 | 0.026 | 0.118 | 0.019 |
| 0.205 | 0.035 | 0.079 | 0.010 |
| 0.163 | 0.023 | 0.121 | 0.024 |
| 0.371 | 0.170 | 0.110 | 0.011 |
| | | 0.148 | 0.014 |
| | | 0.065 | 0.010 |

#### 5.2 um-CpG in colon

**Table S6.** A table containing the normalized fluorescent signal intensity (epigenetic intensity/DNA intensity obtained from the slide scanner)

| Healthy colon (N=18) | | Colon tumor (N=18) | |
|---|---|---|---|
| Average of three to five replicates | Standard deviation | Average of three to five replicates | Standard deviation |
| 0.358 | 0.101 | 0.850 | 0.257 |
| 0.801 | 0.230 | 1.192 | 0.337 |
| 1.057 | 0.065 | 0.692 | 0.087 |
| 0.858 | 0.300 | 0.426 | 0.084 |
| 0.450 | 0.096 | 0.848 | 0.154 |
| 0.798 | 0.179 | 0.151 | 0.027 |
| 0.369 | 0.075 | 0.216 | 0.027 |
| 0.759 | 0.108 | 0.315 | 0.054 |

| | | | |
|---|---|---|---|
| 0.533 | 0.083 | 1.084 | 0.204 |
| 0.309 | 0.067 | 0.431 | 0.200 |
| 0.486 | 0.118 | 0.625 | 0.166 |
| 0.475 | 0.073 | 0.675 | 0.106 |
| 0.215 | 0.030 | 0.452 | 0.173 |
| 0.368 | 0.104 | 0.377 | 0.069 |
| 0.359 | 0.051 | 0.422 | 0.127 |
| 0.121 | 0.059 | 0.384 | 0.042 |
| 0.176 | 0.052 | 0.572 | 0.099 |
| 0.269 | 0.078 | 0.257 | 0.058 |

### 6 Fluorescent signal intensity in hematological samples

#### 6.1 5hmC in hematological samples

**Table S7.** A table containing the normalized fluorescent signal intensity (epigenetic intensity/DNA intensity obtained from the slide scanner)

| Healthy blood (N=18) | | Healthy PBMC (N=14) | |
|---|---|---|---|
| Average of three to five replicates | Standard deviation | Average of three to five replicates | Standard deviation |
| 0.051 | 0.005 | 0.093 | 0.016 |
| 0.052 | 0.025 | 0.048 | 0.004 |
| 0.054 | 0.002 | 0.063 | 0.026 |
| 0.043 | 0.011 | 0.086 | 0.013 |
| 0.052 | 0.003 | 0.081 | 0.022 |
| 0.061 | 0.006 | 0.075 | 0.008 |
| 0.068 | 0.049 | 0.072 | 0.022 |
| 0.106 | 0.082 | 0.089 | 0.019 |
| 0.067 | 0.023 | 0.114 | 0.016 |
| 0.066 | 0.009 | 0.136 | 0.018 |
| 0.037 | 0.016 | 0.079 | 0.029 |
| 0.046 | 0.017 | 0.094 | 0.004 |
| 0.028 | 0.006 | 0.149 | 0.064 |
| 0.038 | 0.017 | 0.128 | 0.063 |
| 0.063 | 0.004 | | |
| 0.075 | 0.015 | | |
| 0.100 | 0.025 | | |
| 0.074 | 0.021 | | |
| **CLL blood (N=18)** | | **CLL PBMC (N=13)** | |
| Average of three to five replicates | Standard deviation | Average of three to five replicates | Standard deviation |
| 0.041 | 0.009 | 0.047 | 0.006 |
| 0.029 | 0.003 | 0.062 | 0.019 |
| 0.031 | 0.007 | 0.053 | 0.006 |
| 0.047 | 0.001 | 0.030 | 0.010 |

| | | | |
|---|---|---|---|
| 0.053 | 0.017 | 0.039 | 0.006 |
| 0.048 | 0.009 | 0.061 | 0.007 |
| 0.037 | 0.010 | 0.037 | 0.012 |
| 0.036 | 0.013 | 0.051 | 0.011 |
| 0.028 | 0.013 | 0.033 | 0.007 |
| 0.035 | 0.005 | 0.042 | 0.005 |
| 0.041 | 0.005 | 0.048 | 0.006 |
| 0.067 | 0.008 | 0.034 | 0.008 |
| 0.052 | 0.007 | 0.049 | 0.021 |
| 0.054 | 0.007 | | |
| 0.046 | 0.014 | | |
| 0.032 | 0.007 | | |
| 0.024 | 0.006 | | |
| 0.038 | 0.008 | | |

#### 6.2 um-CpG in hematological samples

**Table S8.** A table containing the normalized fluorescent signal intensity (epigenetic intensity/DNA intensity obtained from the slide scanner)

| Healthy blood (N=9) | | Healthy PBMC (N=7) | |
|---|---|---|---|
| Average of three to five replicates | Standard deviation | Average of three to five replicates | Standard deviation |
| 0.89 | 0.17 | 0.529 | 0.172 |
| 0.97 | 0.16 | 0.813 | 0.105 |
| 1.15 | 0.91 | 0.753 | 0.415 |
| 0.52 | 0.16 | 1.2840 | 0.3068 |
| 0.65 | 0.11 | 0.6662 | 0.0972 |
| 1.02 | 0.24 | 0.4106 | 0.0056 |
| 0.69 | 0.10 | 0.5800 | 0.1594 |
| 0.11 | 0.03 | | |
| 0.53 | 0.31 | | |
| **CLL blood (N=6)** | | **CLL PBMC (N=10)** | |
| Average of three to five replicates | Standard deviation | Average of three to five replicates | Standard deviation |
| 0.78 | 0.09 | 0.95 | 0.13 |
| 0.85 | 0.26 | 0.82 | 0.20 |
| 1.47 | 0.58 | 1.45 | 0.38 |
| 2.75 | 0.90 | 1.70 | 0.22 |
| 2.00 | 0.51 | 1.23 | 0.28 |
| 1.42 | 0.40 | 3.18 | 0.43 |
| | | 0.70 | 0.16 |
| | | 0.84 | 0.27 |
| | | 1.18 | 0.69 |
| | | 1.00 | 0.25 |
